# Pich, an ATP-dependent chromatin remodeling protein, transcriptionally co-regulates oxidative stress response

**DOI:** 10.1101/2021.07.21.453306

**Authors:** Anindita Dutta, Apurba Das, Deepa Bisht, Vijendra Arya, Rohini Muthuswami

## Abstract

Cells respond to oxidative stress by elevating the levels of antioxidants, signaling, and transcriptional regulation often implemented by chromatin remodeling proteins. The study presented in this paper shows that the expression of PICH, an ATP-dependent chromatin remodeler, is upregulated during oxidative stress in HeLa cells. We also show that PICH regulates the expression of Nrf2, a transcription factor regulating antioxidant response, both in the absence and presence of oxidative stress. In turn, Nrf2 regulates the expression of PICH in the presence of oxidative stress. Both PICH and Nrf2 together regulate the expression of antioxidant genes and this transcriptional regulation is dependent on the ATPase activity of PICH. In addition, H3K27ac modification also plays a role in activating transcription in the presence of oxidative stress. Co-immunoprecipitation experiments show that PICH and Nrf2 interact with H3K27ac in the presence of oxidative stress. Mechanistically, PICH recognizes ARE sequences present on its target genes and introduces a conformational change to the DNA sequences leading us to hypothesize that PICH regulates transcription by remodeling DNA. PICH ablation leads to reduced expression of Nrf2 and impaired antioxidant response leading to increased ROS content, thus, showing PICH is essential for the cell to respond to oxidative stress.

## INTRODUCTION

The ATP-dependent chromatin remodeling proteins dynamically alter the chromatin architecture to enable transcription factors to access the genomic DNA and thereby mediate transcription and repair [1]. Organismal life encounters reactive oxidants from internal metabolism as well as from environmental toxicant exposure resulting in oxidative stress due to imbalance between reactive oxygen species (ROS) and antioxidants. The role of epigenetic modulators in combating oxidative stress has been well-documented. For example, oxidative stress has been shown to activate histone acetyltransferases and inhibit histone deacetylase activity [2]. Studies have also shown that BRG1, an ATP-dependent chromatin remodeling protein, interacts with Nrf2, a transcription factor, to regulate the expression of *HO-1* in response to oxidative stress [3]. CHD6 too is an important regulator of oxidative stress [4].

PICH (PLK-1 interacting checkpoint helicase), also known as ERCC6L, is a Rad54-like helicase belonging to the ATP-dependent chromatin remodeling protein family [5–7]. The protein has been identified as a strong binding partner and substrate of PLK-1 that localizes at the kinetochores [5]. Immunofluorescence staining has revealed that PICH is mostly concentrated in between kinetochores in prometaphase cells while in metaphase cells the protein localizes to numerous short threads that stretch between the sister-kinetochores of the aligned chromosomes [5]. The protein has been shown to respond to the tension-dependent alterations in DNA topology by resolving the ultrafine bridges generated in between the centromeres of the segregating chromatids and regulates the spindle attachment checkpoint [8]. PICH depleted cells show chromosomal abnormalities, dead cells, bi-nuclei, and multi-nuclei formation [9]. The protein is found to be overexpressed in many cancers and silencing the expression of PICH has shown to inhibit their proliferation [10, 11]. Studies have also shown that the protein interacts with BEND3 as well as with topoisomerases; however, the protein has not been shown to reposition/remodel nucleosomes [9,12,13].

The nuclear factor erythroid 2 (NFE2)-related factor 2 (Nrf2) is a member of the cap ‘n’collar (CNC) subfamily of basic region leucine zipper transcription factors [14, 15]. Studies over the past decade have established the role of Nrf2 in combating oxidative stress in cells [16]. In cells, in the absence of oxidative stress, Nrf2is ubiquitinated by Keap1 and degraded by the proteasome pathway [17, 18]. In the presence of oxidative stress, Keap1 is inactivated, leading to the release of Nrf2 which migrates to the nucleus and activates the expression of antioxidant genes by binding to the Antioxidant Responsive Element (ARE) present on the promoters of the antioxidant genes [17, 18]. Knockout of *Nrf2* in mice increases their susceptibility to chemical toxins and the mice exhibit disease conditions associated with oxidative pathology [19].

In this paper, the role of PICH during oxidative stress has been investigated. We provide evidence that PICH regulates the antioxidant response in HeLa cells under oxidative stress. PICH also regulates the expression of *Nrf2* and together with Nrf2 and H3K27ac appears to drive the expression of the antioxidant genes. We also show that PICH induces a conformational change in the DNA structure in an ATP-dependent manner, thus, provide a possible mechanism for transcriptional co-regulation mediated by this protein.

## RESULTS

### PICH expression is upregulated when cells are exposed to oxidative stress

Previously, we reported that BRG1 and SMARCAL1 both belonging to the ATP-dependent chromatin remodeling protein family co-regulate each other’s expression on treatment with doxorubicin [20]. To investigate whether this regulation is universal and occurs on induction of any type of DNA damage, HeLa cells were subjected to oxidative stress generated endogenously by transfecting cells with a plasmid encoding for D-amino acid oxidase (DAAO). The cells, 36 h post-transfection, were treated with 25 mM D-serine for 10 min [21] resulting in the production of H_2_O_2_ within the cell by the action of DAAO on D-serine. ROS production was confirmed by DCFDA staining (Fig. S1A and B). The treated cells were harvested, and the RNA was isolated to analyze the expression of *BRG1* and *SMARCAL1* by qPCR. Surprisingly, the expression of *BRG1* and *SMARCAL1* were unchanged but the expression of *PICH*, an ATP-dependent chromatin remodeler, and *Nrf2*, a transcription factor, was upregulated (Fig S1C). The changes in the transcript level were confirmed by western blot (Fig. S1D and E). Further, the expression of antioxidant genes *CAT, GPX1, GSR,* and *TXNRD1*was found to be upregulated while SOD1 was unchanged (Fig. S1F). Concomitantly, catalase activity was also found to be upregulated (Fig S1G). Untransfected HeLa cells treated with D-serine did not exhibit these changes in the transcript indicating that this molecule by itself is not contributing to oxidative stress or changes thereof (Fig. S1H). Taken together, the experimental results indicated that PICH expression was upregulated on oxidative stress.

To confirm the upregulation of PICH in response to oxidative stress, HeLa cells were treated with 100 µM H_2_O_2_ and the expression was analyzed as a function of time [22]. Analysis showed that the expression of *PICH* was upregulated 20 min post-treatment in comparison to the untreated HeLa cells (Fig. 1A). The upregulation was confirmed by western blot (Fig. 1B and C). As the first peak of *PICH* expression was at 20 min post-treatment as compared to the untreated HeLa cells, all the experiments described henceforth were performed after treating HeLa cells with 100 μM H_2_O_2_ for 20 min. Under this condition, the production of ROS was confirmed by staining the cells with DCFDA (Fig. 1D and E). γH2AX foci formation confirmed DNA damage was also induced under this condition (Fig. 1F and G).

**Figure 1.**
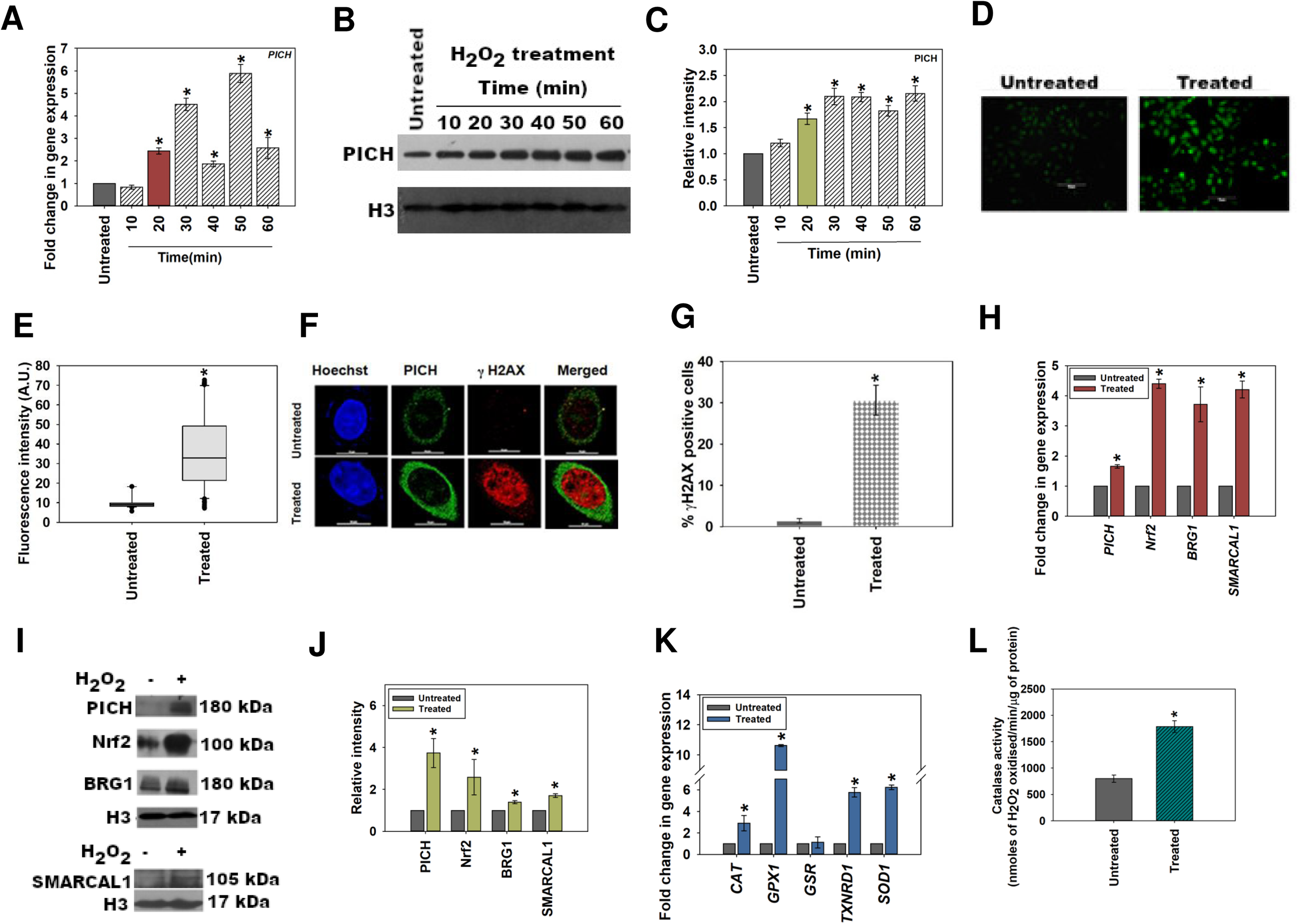
PICH expression is upregulated when cells are exposed to oxidative stress. (A). Expression of *PICH* was analyzed at indicated time points in HeLa cells after treatment with 100 µM H_2_O_2_ by qPCR. (B). Expression of PICH was examined by western blot. H3 was used as an internal control. (C). Quantitation of the western blot using Image J software. (D). Cellular ROS was analyzed using DCFDA in HeLa cells in the absence and presence of H_2_O_2_. (E). Fluorescent intensity was quantitated using the software provided by TiE, Nikon Microscope. (F). gH2AX foci formation was visualized absence and presence of H_2_O_2_ in HeLa cells using confocal microscope. (G). Quantitation of the gH2AX positive cells. (H). Expression of *PICH*, *Nrf2, BRG1*, and *SMARCAL1* were quantitated using qPCR. (I). Expression of PICH, Nrf2, BRG1, and SMARCAL1 was analyzed by western blot. (J). Quantitation of the western blot using ImageJ software. (K). Expression of antioxidant genes *CAT*, *GPX1*, *GSR*, *TXNRD1,* and *SOD1* was analyzed by qPCR. (L). Catalase activity (μmol/min) was quantitated in untreated and treated (100 mM H_2_O_2_; 20 min) HeLa cells. *GAPDH* was used as the internal control in all the qPCR experiments. The qPCR experiments are presented as average ± SEM of three independent experiments. The star indicates p-value is < 0.05. The western blots are presented as average ± SEM of two individual experiments. The intensities of the proteins were normalized with respect to H3 and are represented as Relative Intensity in the y-axis.

Next, the expression of *Nrf2, BRG1*, and *SMARCAL1* was analyzed after treatment with 100 μM H_2_O_2_ for 20 min. As expected, *Nrf2* expression was upregulated on the generation of oxidative stress exogenously (Fig. 1H). Interestingly, unlike the endogenous pathway, on exogenous treatment with H_2_O_2_, *BRG1* and *SMARCAL1* were upregulated along with *Nrf2* (Fig. 1H). This upregulation was also confirmed by western blots (Fig. 1I and J).

The dichotomy between the endogenous and exogenous pathways was further accentuated by analysis of the expression of antioxidant genes. Unlike the endogenous pathway, where *SOD1* was unchanged and *GSR* was upregulated, in the exogenous pathway, *SOD1* was upregulated while *GSR* was unchanged (compare Fig. 1K with Fig. S1F). As it is not possible to compare the amount of ROS produced under these two conditions, therefore, a direct comparison with the amount of ROS produced and the pathway activated cannot be made. We, however, noted that catalase activity was upregulated when cells were treated with D-serine after *DAAO* transfection as well as with hydrogen peroxide (Fig. 1L and Fig. S1G).

### PICH expression on oxidative stress is cell line dependent

Various reports have suggested that the expression of a given chromatin remodeling protein in response to stress is cell line-specific [3, 23]. To understand whether *PICH* expression in response to oxidative stress was also dependent on the cell line, transcript levels were analyzed in SiHa and HEK293 cell lines in response to 100 μM H_2_O_2_ treatment given for 20 min. In HEK293 cells, *PICH* and *Nrf2* expression were upregulated like in HeLa cells while in SiHa cells, *PICH* transcript level was downregulated and *Nrf2* was upregulated (Fig. S1I and J). Thus, the expression of PICH in response to oxidative stress does appear to be cell line-specific.

In this paper, the connection between PICH, Nrf2, and antioxidant genes on treatment with H_2_O_2_ in HeLa cells has been explored.

### PICH regulates the expression of Nrf2 in HeLa cells in the absence and presence of oxidative stress

To understand whether PICH is necessary for the upregulation of *Nrf2*, on oxidative stress, shRNA against the 3′ UTR of PICH was used to downregulate the expression of this gene in HeLa cells. Downregulation of *PICH* led to downregulation of *Nrf2* both in the absence (P-value = 3E-10) and presence (P-value =2.21E-06) of H_2_O_2_, indicating PICH regulates *Nrf2* expression regardless of oxidative stress (Fig. S2A and Fig. 2A). The downregulation was confirmed by western blots (Fig. S2B-C and Fig. 2B-C). Further, the expression of antioxidant genes was also downregulated as was catalase activity both in the absence and presence of oxidative stress (Fig. S2D-E and Fig. 2D-E respectively). ROS levels, as analyzed by the DCFDA label, were found to be significantly upregulated (P-value = 0.00) in Sh*PICH* cells as compared to HeLa cells transfected with scrambled shRNA both in the absence and presence of H_2_O_2_ (Fig. S2F-G and Fig. 2F-G respectively). PICH downregulation has been reported to increase DNA damage [24]. Increased number of γH2AX foci positive cells in Sh*PICH* cells as compared to ScrRNA cells were observed both in the absence (P = 0.01) and presence (P = 8.04E-08) of H_2_O_2_ (Fig. S2H-I and Fig. 2H-I). Furthermore, the number of γH2AX foci positive Sh*PICH* cells were more in the treated condition as compared to untreated condition (P-value = 0.006) indicating increased DNA damage in the presence of oxidative stress when PICH expression is ablated.

**Figure 2.**
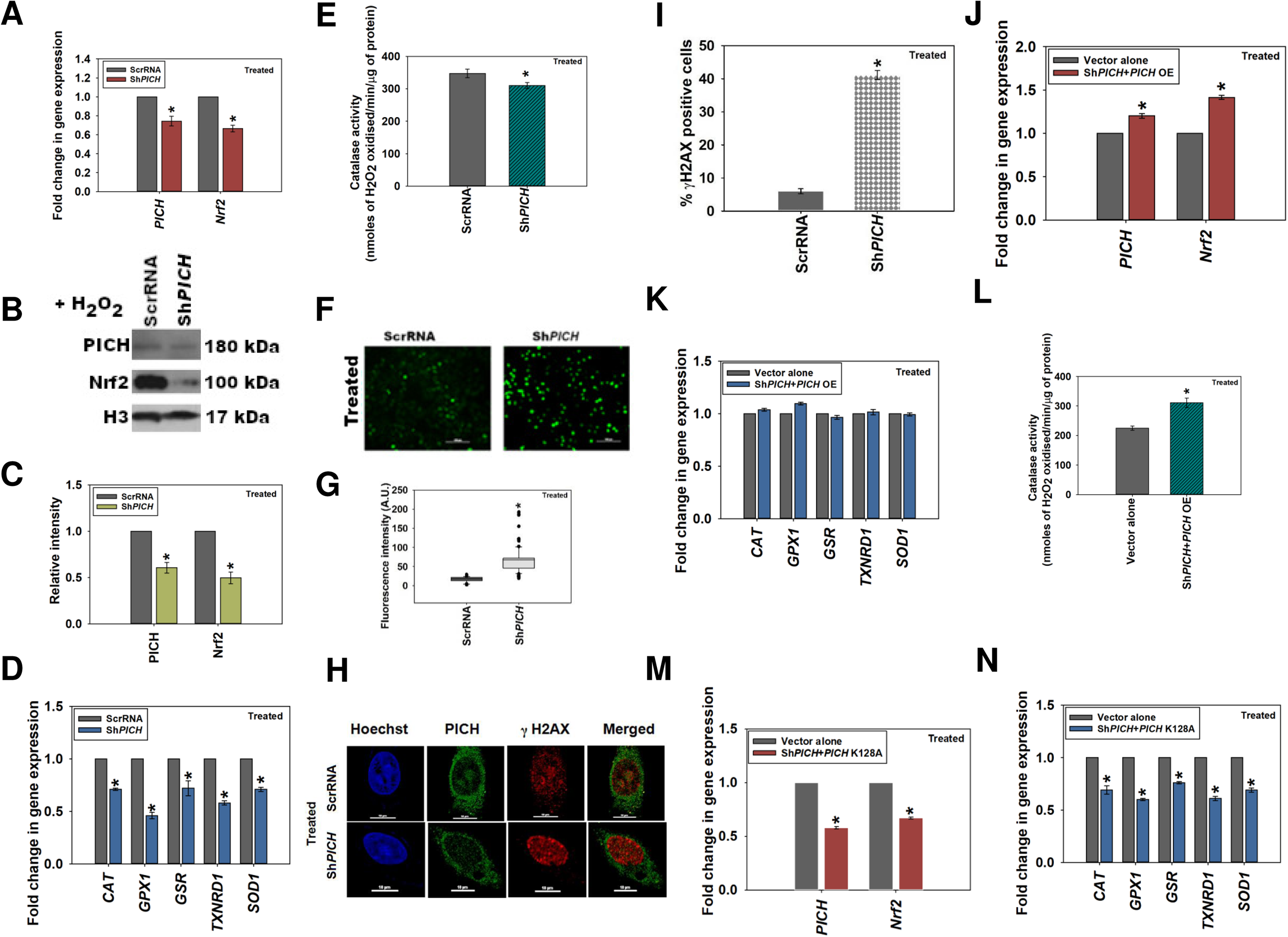
PICH regulates the expression of *Nrf2* in HeLa cells in the absence and presence of oxidative stress. (A). Expression of *PICH* and *Nrf2* in HeLa cells transfected with either ScrRNA or with Sh*PICH* plasmid in HeLa cells after treatment with 100 μM H_2_O_2_ for 20 min. (B). Expression of PICH and Nrf2 in treated (100 μM H_2_O_2_; 20 min) HeLa cells transfected with either ScrRNA or with Sh*PICH* was analyzed by western blot. (C). Quantitation of the western blots. (D). Expression of antioxidant genes *CAT*, *GPX1*, *GSR*, *TXNRD1*, and *SOD1* were quantitated by qPCR in treated (100 μM H_2_O_2_; 20 min) HeLa cells transfected either with ScrRNA or with Sh*PICH*. (E). Catalase activity (μmol/min) was quantitated in treated (100 μM H_2_O_2_; 20 min) HeLa cells transfected either with ScrRNA or with Sh*PICH*. (F). Cellular ROS was analyzed using DCFDA in treated HeLa cells transfected either with ScrRNA or with Sh*PICH*. (G). Fluorescent intensity was quantitated using the software provided by TiE, Nikon Microscope. (H). gH2AX foci formation was visualized in treated HeLa cells transfected either ScrRNA or with ShPICH using confocal microscope. (I). Quantitation of the gH2AX positive cells. (J). Transcript levels of *PICH* and *Nrf2* were quantitated by qPCR in treated (100 μM H_2_O_2_; 20 min) HeLa cells transfected with ScrRNA and empty vector or with the Sh*PICH* and *PICH* overexpression construct. (K). Expression of antioxidant genes *CAT*, *GPX1*, *GSR*, *TXNRD1*, and *SOD1* were quantitated in treated (100 μM H_2_O_2_; 20 min) HeLa cells transfected with ScrRNA and empty vector or with the Sh*PICH* and *PICH* overexpression construct. (L). Catalase activity (μmol/min) was estimated in treated (100 mM H_2_O_2_; 20 min) HeLa cells transfected with ScrRNA and empty vector or with the Sh*PICH* and *PICH* overexpression construct. (M). Transcript levels of *PICH* and *Nrf2* were quantitated by qPCR in treated (100 μM H_2_O_2_; 20 min) HeLa cells transfected with ScrRNA and empty vector or with the Sh*PICH* and *PICH* K128A overexpression construct. (N). Expression of antioxidant genes *CAT*, *GPX1*, *GSR*, *TXNRD1*, and *SOD1* were quantitated in treated (100 μM H_2_O_2_; 20 min) HeLa cells transfected with ScrRNA and empty vector or with the Sh*PICH* and *PICH* K128A overexpression construct. *GAPDH* was used as the internal control in the qPCR experiments. The qPCR experiments are presented as average ± SEM of three independent experiments. The western blots are presented as average ± SEM of three independent experiments. The intensities were quantitated using Image J software. The intensities of the proteins were normalized with respect to H3 and are represented as Relative Intensity in the y-axis. The knockdown efficiency of *PICH* was 50-60% in all the experiments performed.

The downregulation of *NRF2* could be rescued by overexpressing *PICH* in *PICH* downregulated cells in both untreated and treated HeLa cells (Fig. S2J and Fig. 2J). The expression of antioxidant genes, as well as catalase activity, was also found to be rescued in H_2_O_2_ treated cells co-transfected with Sh*PICH* along with *PICH* overexpression cassette as compared to cells transfected with vector alone (Fig. 2K and L; untreated controls are shown in Fig. S2K and L).

### The ATPase activity of PICH is required for transcriptional regulation

To understand whether the ATPase activity of the protein is required for transcriptional regulation, the invariant K present in the conserved helicase motif I (GLGK^128^T) was mutated to alanine in the full-length PICH [9]. Co-transfection of the plasmid expressing the mutant PICH (PICHK128A) with Sh*PICH* into HeLa cells showed that the expression of Nrf2, as well as antioxidant genes, was not rescued when transfected cells were treated 100 μM H_2_O_2_ for 20 min highlighting the importance of ATPase activity of PICH in transcriptional regulation (Fig. 2M and N; untreated controls are shown in Fig. S2M and N).

Taken together, PICH regulates the expression of *Nrf2* both in the absence and presence of oxidative stress. Further, PICH also regulates the genes encoding for antioxidants. Finally, the ATPase activity of the protein is required for transcriptional regulation as overexpression of the ATPase dead mutant failed to rescue the downregulation of genes in Sh*PICH* cells.

### Nrf2 regulates the expression of PICH in HeLa cells in the presence of oxidative stress

As the expression of PICH was upregulated on oxidative stress, we asked whether Nrf2 has a role in regulating the expression of this gene on oxidative stress. Nrf2 is a transcription factor that binds to Antioxidant Response Element (ARE) and thus, regulates the expression of antioxidant genes [25]. The polymorphic ARE core sequence 5′-RTGYCNNNGCR-3′, where R is a purine and Y is a pyrimidine, is present on the promoters of genes encoding for antioxidant genes [26, 27]. An *in silico* analysis identified ARE sequence was present on the *PICH* promoter leading us to hypothesize that Nrf2 could potentially regulate the expression of this gene. (Supplementary Table 3).

*Nrf2* expression was downregulated by transiently transfecting HeLa cells with shRNA against the 3′ UTR of the gene. Downregulation of *Nrf2* led to reduced expression of *PICH* both in the absence (P-value = 3.56E-06) and presence (P-value = 1.65E-05) of oxidative stress (Fig. 3A-F). The expression of PICH in *Nrf2* downregulated cells could be rescued by overexpressing *Nrf2* in the presence of oxidative stress (Fig. 3G and H; heat maps shown in Fig. S3)

**Figure 3.**
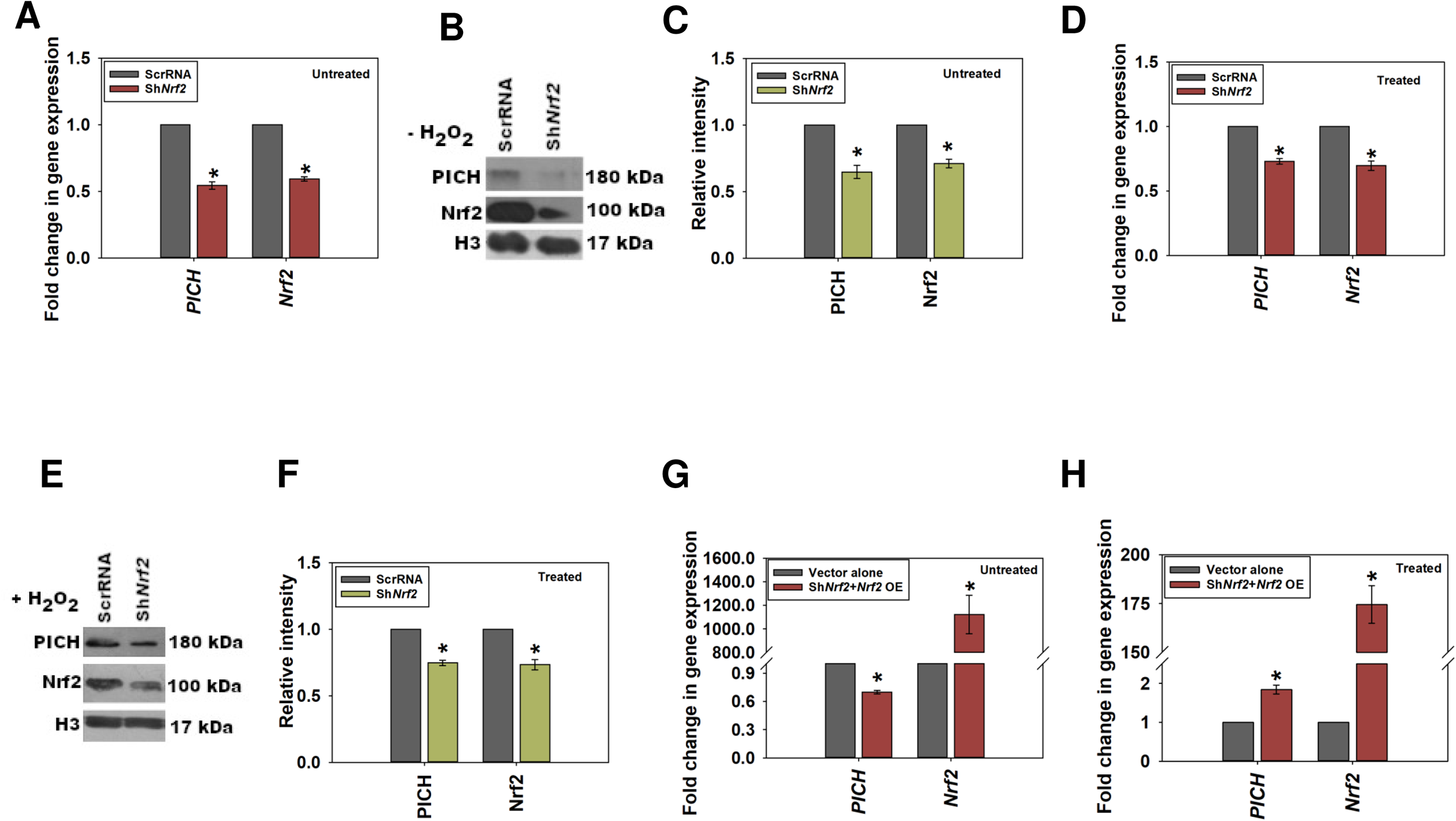
Nrf2 regulates the expression of *PICH* in HeLa cells in the presence of oxidative stress. (A). Expression of *PICH* and *Nrf2* in untreated HeLa cells transfected either with ScrRNA or with Sh*Nrf2* construct. (B). Expression of PICH and Nrf2 was analyzed by western blot in untreated HeLa cells transfected either with ScrRNA or with the Sh*Nrf2* construct. (C). Quantitation of the western blots. (D). Expression of *PICH* and *Nrf2* in treated (100 μM H_2_O_2_; 20 min) HeLa cells transfected either with ScrRNA or with Sh*Nrf2* construct. (E). Expression of PICH and Nrf2 was analyzed by western blot in treated (100 μM H_2_O_2_; 20 min) HeLa cells transfected either with ScrRNA or with Sh*Nrf2* construct. (F) Quantitation of the western blots. (G). Transcript levels of *PICH* and *Nrf2* were estimated by qPCR in untreated HeLa cells transfected either with ScrRNA and empty vector or with Sh*Nrf2* construct along with *Nrf2* overexpression plasmid constructs. (H). Transcript levels of *PICH* and *Nrf2* were estimated by qPCR in treated (100 μM H_2_O_2_; 20 min) HeLa cells transfected either with ScrRNA and empty vector or with Sh*Nrf2* construct along with *Nrf2* overexpression plasmid constructs. *GAPDH* was used as the internal control in the qPCR experiments. The qPCR experiments are presented as average ± SEM of three independent experiments. The western blots are presented as average ± SEM of three independent experiments. The intensities were quantitated using Image J software. The intensities of the proteins were normalized with respect to H3 and are represented as Relative Intensity in the y-axis. The knockdown efficiency of *Nrf2* was 35-45% in all the experiments performed.

From these experimental results, we concluded that in HeLa cells, Nrf2 regulates the expression of *PICH* only on oxidative stress.

Thus, PICH regulates *Nrf2* both in the absence and presence of oxidative stress while Nrf2 regulates *PICH* only in the presence of oxidative stress.

### The occupancy of PICH and Nrf2 increases on the promoters of effector genes on oxidative stress

To understand the mechanism of transcriptional regulation by PICH, ChIP experiments were done to probe the occupancy of PICH, Nrf2, and RNAPII on *PICH, Nrf2, SOD1, GPX1* and *TXNRD1* promoters in the absence and presence of oxidative stress.

On the *PICH* promoter, PICH, Nrf2, and RNAPII occupancy increased only on oxidative stress (Fig. 4A; heat map shown in Fig. S4B). This suggests that Nrf2 does regulate the expression of *PICH* on oxidative stress. Further, PICH appears to co-regulate its own transcription.

**Figure 4.**
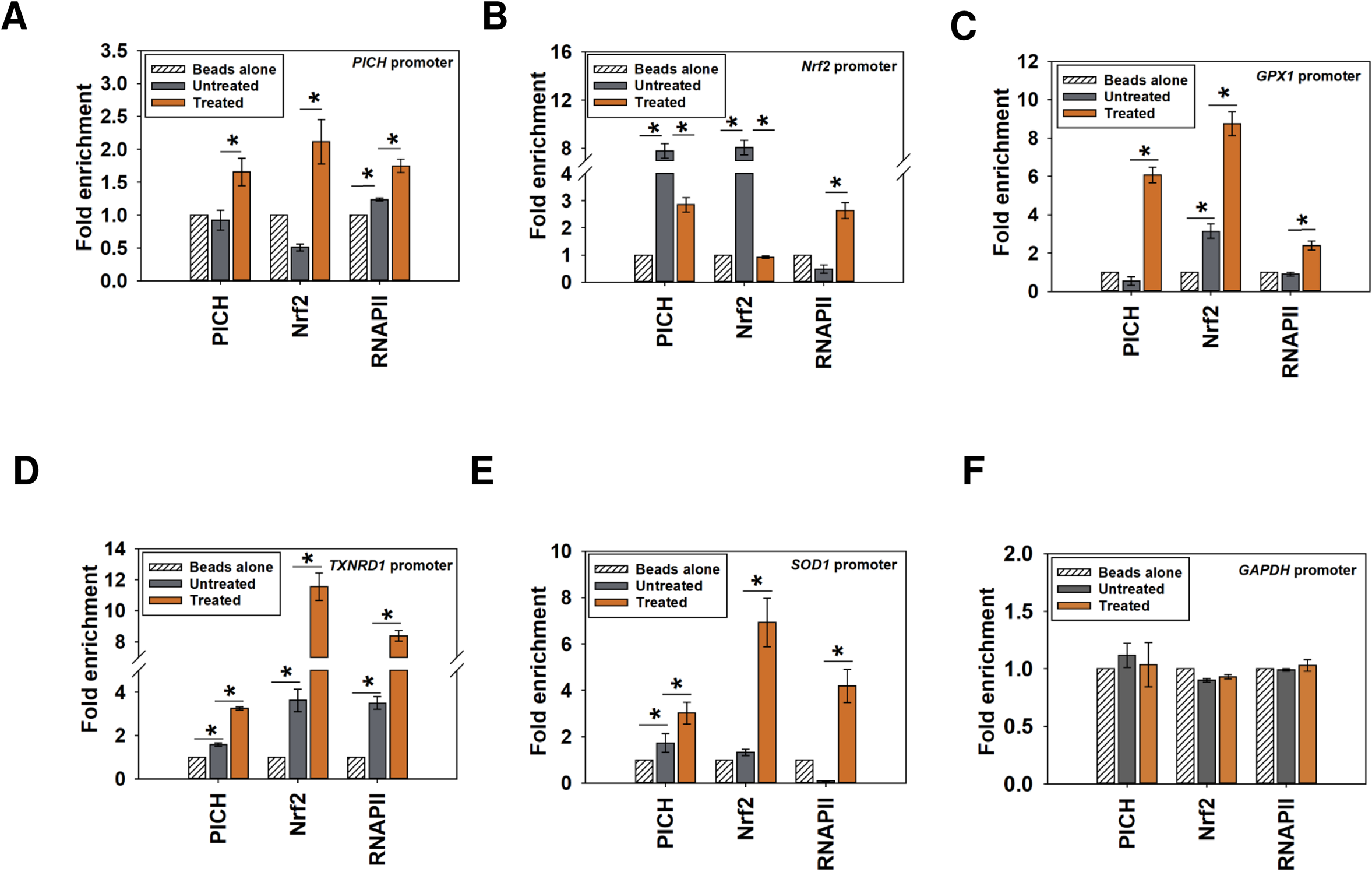
The occupancy of PICH and Nrf2 increases on the promoters of effector genes on oxidative stress. The occupancy of PICH, Nrf2, RNAPII on the (A). *PICH* promoter; (B). *Nrf2* promoter; (C). *GPX1*promoter; (D). *TXNRD1* promoter; (E). *SOD1* promoter; and (F). *GAPDH* promoter was analyzed by ChIP in untreated and treated (100µM H_2_O_2_ for 20 min) HeLa cells. In these experiments, primers were made to probe a region −300 bp upstream and +200 bp downstream of the transcription start site (TSS). *GAPDH* promoter was used as the negative control. The ChIP data are presented as average ± SEM of three independent experiments. Star indicates significance at p-value is<0.05.

PICH and Nrf2 were both found to be present on the *Nrf2* promoter; their occupancy decreased with oxidative stress as compared to the occupancy in the absence of oxidative stress (Fig. 4B; heat map shown in Fig. S4B). Overall, PICH regulates Nrf2 both in the absence and presence of oxidative stress. RNAPII occupancy, on the other hand, increased on the promoter on oxidative stress correlating with increased transcription of *Nrf2* (Fig. 4B; heat map shown in Fig. S4B).

Finally, PICH, Nrf2, and RNAPII occupancy increased on *GPX1*, *TXNRD1,* and *SOD1* promoters on oxidative stress, correlating with increased transcription (Fig. 4C-E; heat map shown in Fig. S4B).

In contrast, the occupancy of PICH, Nrf2, and RNAPII did not alter on *GAPDH* promoter (Fig. 4F; heat map shown in Fig. S4B) indicating that the change in occupancy was specific to the genes involved in oxidative stress response.

Thus, PICH binds to the promoter (ARE) sequences of its target genes and co-regulates transcription with Nrf2.

### Histone marks associated with transcription activation are enriched on PICH, Nrf2, and antioxidant gene promoters on oxidative stress

We hypothesized that the increased transcription of *PICH*, *Nrf2,* and antioxidant genes should be accompanied by an increase in activation histone marks. H3K9ac, H3K4me2, H3K4me3, and H3K27ac modifications are associated with transcription activation [28–30].

Analysis of the ChIP-seq data of these histone marks-H3K9c (GSM733756), H3K4me2 (GSM733734), H3K27ac (GSM733684), and H3K4me3 (GSM733682)-available in the public domain [28] showed that these are present on the promoters of *PICH, Nrf2, GPX1, TXNRD1* and *SOD1* in untreated HeLa cells (Fig. S4A).

Next, ChIP experiments were done to understand the enrichment of H3K4me3 and H3K27ac on *PICH*, *Nrf2*, *GPX1*, *TXNRD1*, and *SOD1* promoters in the absence and presence of oxidative stress.

H3K4me3 and H3K27ac modifications were enriched on *PICH* promoter in the treated cells as compared to untreated cells (Fig. 5A). On the *Nrf2* promoter, however, enrichment of only H3K27ac was observed in the treated cells as compared to the untreated cells indicating that only this modification is playing a role in transcriptional regulation of this gene (Fig. 5B). Finally, both H3K4me3 and H3K27ac modifications were found to be enriched on *GPX1*, *TXNRD1*, and *SOD1* promoters in the presence of oxidative stress, indicating both these modifications are associated with transcriptional activation of these genes (Fig. 5C-E). In contrast, neither of these modifications were enriched on the GAPDH promoter (Fig. 5F).

**Figure 5.**
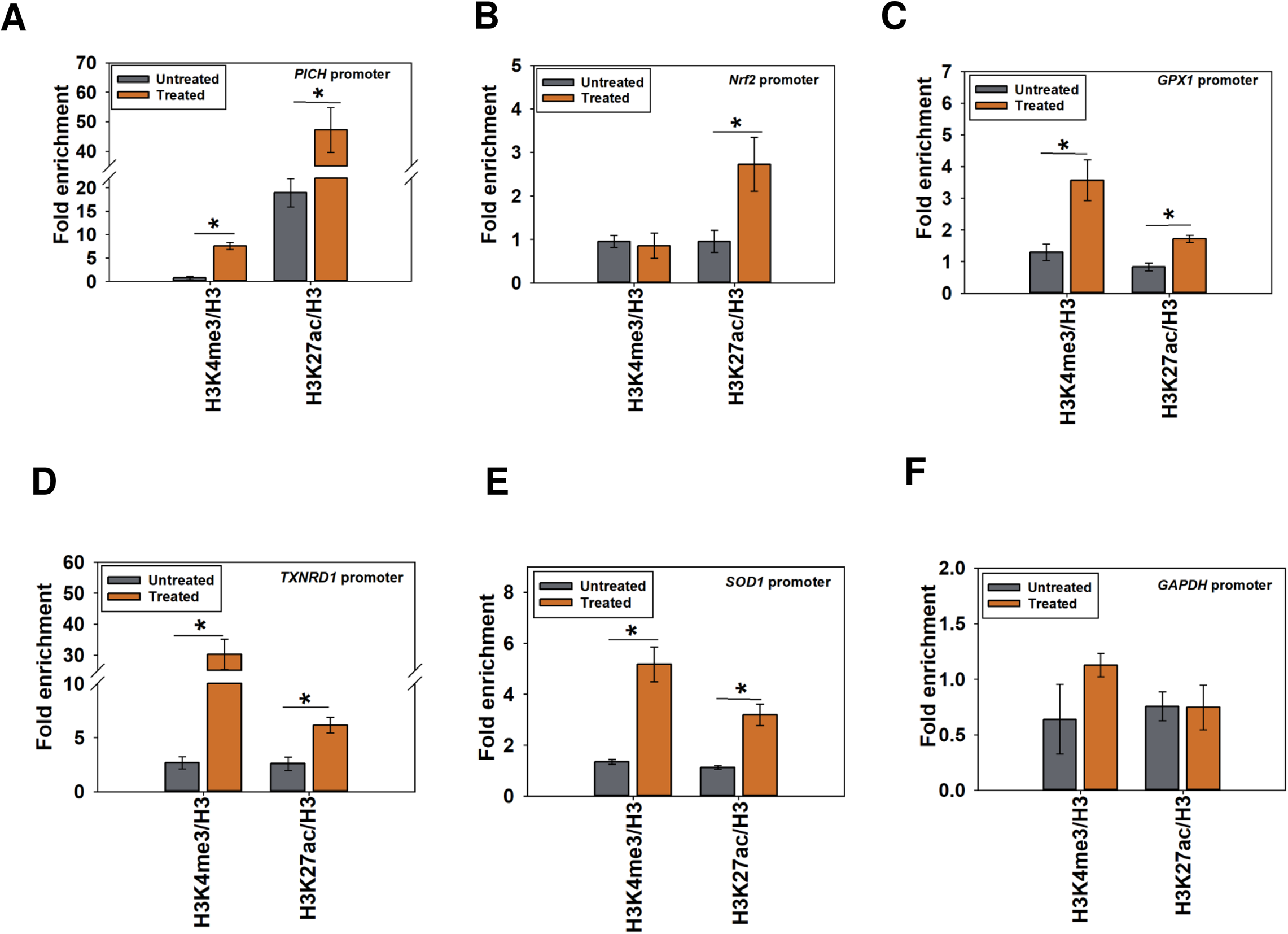
Histone marks associated with transcription activation are enriched on PICH, Nrf2, and antioxidant gene promoters on oxidative stress. The fold enrichment of H3K4me3 and H3K27ac as a ratio of H3 was determined using ChIP on the (A). *PICH* promoter; (B). *Nrf2* promoter; (C). *GPX1* promoter; (D). *TXNRD1* promoter; (E). *SOD1* promoter; and (F) *GAPDH* promoter in untreated and treated (100 μM H_2_O_2;_ 20 min) HeLa cells. In these experiments, primers were made to probe a region −300 bp upstream and +200 bp downstream of the transcription start site (TSS). *GAPDH* promoter was used as the negative control. The ChIP data are presented as average ± SEM of three independent experiments. Star indicates significance at p-value is<0.05.

Thus, from these experiments, it was concluded that both H3K4me3 and H3K27ac were associated with transcriptional regulation of *PICH* and antioxidant genes on oxidative stress. Further, only H3K27ac was playing a role in transcriptional activation of the *Nrf2* gene on oxidative stress.

### The occupancy of Nrf2, RNAPII, and H3K27ac on the target genes is dependent on PICH expression

Next, we investigated whether PICH is needed for the recruitment of Nrf2, RNAPII, and H3K27ac to the promoters of the target genes on oxidative stress. ChIP experiments were performed using cells transfected with either scrambled shRNA (ScrRNA) or shRNA against *PICH* (Sh*PICH*) and treated with H_2_O_2_.

The occupancy of RNAPII and H3K27ac decreased on *PICH*, *Nrf2*, *GPX1*, *TXNRD1,* and *SOD1* promoters in the Sh*PICH* cells as compared to ScrRNA cells on oxidative stress correlating with decreased expression of these genes (Fig. 6A-E). Nrf2 occupancy too decreased on *PICH*, *SOD1*, *GPX1*, and *TXNRD1* promoters; however, it increased on its own promoter in Sh*PICH* cells as compared to ScrRNA on oxidative stress (Fig. 6B). In contrast, the occupancy of Nrf2, RNAPII, and H3K27ac did not alter on the *GAPDH* promoter (Fig. 6F).

**Figure 6.**
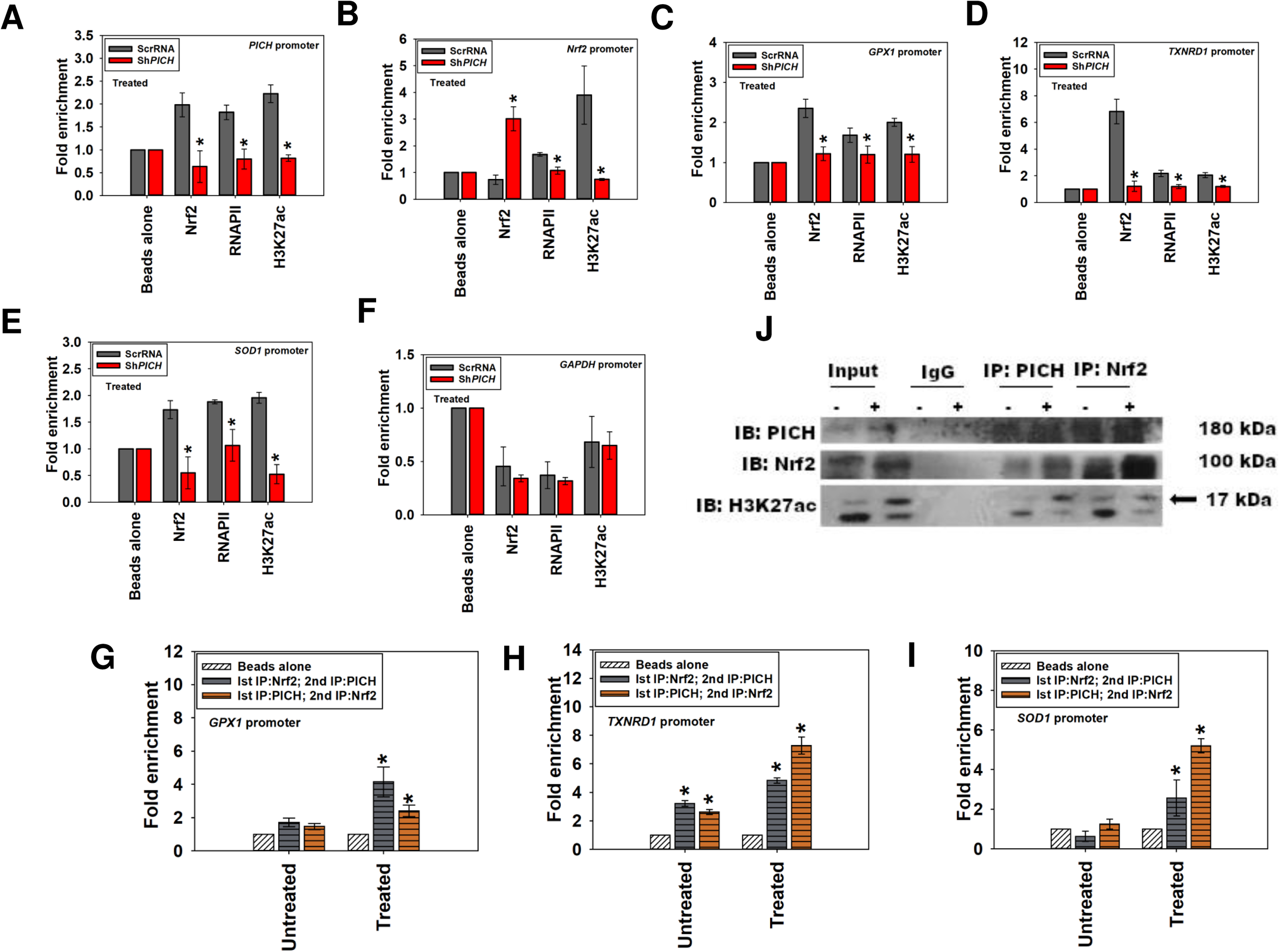
The occupancy of Nrf2, RNAPII, and H3K27ac on the target genes is dependent on *PICH* expression. Occupancy of Nrf2, RNAPII, and H3K27ac was assessed using ChIP in HeLa cells transfected either with ScrRNA or Sh*PICH* after treatment with 100 μM H_2_O_2_ for 20 min on (A). *PICH* promoter; (B). *Nrf2* promoter; (C). *GPX1* promoter; (D). *TXNRD1* promoter; (E). *SOD1* promoter; and (F). *GAPDH* promoter. Simultaneous occupancy of PICH and Nrf2 was assessed in untreated and treated HeLa cells using ChIP re-ChIP on (G). *GPX1* promoter; (H). *TXNRD1* promoter and (I). *SOD1* promoter. (J). Interaction between PICH, Nrf2, and H3K27ac was probed in untreated and treated (100 μM H_2_O_2_; 20 min) HeLa cells by co-immunoprecipitations. IgG was taken as the negative control. In these experiments, primers were made to probe a region −300 bp upstream and +200 bp downstream of the transcription start site (TSS). The ChIP data are presented as average ± SEM of three independent experiments. Star indicates significance at p-value is<0.05. In the ChIP-reChIP experiments, primers were made to probe a region −300 bp upstream and +200 bp downstream of the transcription start site (TSS). The ChIP-reChIP data for *SOD1*, *GPX1*, and *TXNRD1* promoters are presented as average ± SEM of two independent experiments. Star indicates significance at p-value is<0.05.

Taken together, we hypothesize that PICH is needed for the recruitment of transcription machinery to the target genes on oxidative stress.

### PICH interacts with both Nrf2 and H3K27ac modification

Next, we asked whether PICH is present on the target promoters simultaneously with Nrf2. ChIP-reChIP experiments confirmed that PICH and Nrf2 are indeed present together on the promoter regions of *GPX1*, *TXNRD1*, and *SOD1* (Fig. 6G-I).

Co-immunoprecipitation showed that PICH interacted either directly or indirectly with both Nrf2 and H3K27ac (Fig. 6J). This interaction was present both in the absence and presence of oxidative stress. However, the H3K27ac modification appears to increase on treatment with H_2_O_2_. Both anti-PICH antibody and anti-Nrf2 antibody appear to pull down this modification on oxidative stress (Fig. 6J). Immunoprecipitation with anti-Nrf2 antibody also pulled down PICH both in the absence and presence of oxidative stress while it pulled down H3K27ac in the presence of oxidative stress (Fig. 6J).

Thus, from these experiments, we conclude that PICH and Nrf2 are simultaneously present on the promoters of the target genes. Further, PICH and Nrf2 form a complex in the presence of H3K27ac in HeLa cells.

### PICH alters the conformation of DNA in an ATP-dependent manner

Studies have shown that PICH does not reposition nucleosomes [8]. In such a scenario, how does PICH co-regulate transcription? Studies have shown that BRG1, as well as SMARCAL1, can induce conformational changes in DNA [20,31,32]. Therefore, we hypothesized that PICH might be co-regulating transcription by remodeling DNA.

Recombinant PICH (1-750 aa; 86 kDa; hereinafter referred to as ΔPICH) was overexpressed and purified from *E. coli* as explained in the Methods section (Fig. 7A and B). The protein showed maximal DNA-dependent ATPase activity in the presence of stem-loop DNA (slDNA) (Fig. 7C). Mutation of the lysine (K128; hereinafter referred to as ΔPICHK128A) residue of the conserved GKT residue resulted in the loss of DNA-dependent ATPase activity (Fig. S5A and Fig. 7D). Next, the ability of the recombinant PICH to hydrolyze ATP was analyzed in the presence of ARE sequences as ChIP experiments had shown that the protein is bound to these regions. As maximal ATPase activity was observed in the presence of slDNA, we hypothesized that the protein most probably was recognizing specific structural elements present in the DNA. Mfold [33] showed that the sequence where PICH occupancy was observed by ChIP can potentially form stem-loop structures while QGRS mapper [34] showed these sequences have a low potential for the formation of G-quadruplexes (Supplementary Table 3). Therefore, these sequences were denatured by heating to 95°C and then either slow-cooled at room temperature (facilitates inter-strand annealing) or fast-cooled at 4°C (facilitates intra-strand annealing). ATPase assays showed that ΔPICH but not ΔPICHK128A was able to hydrolyze ATP in presence of all the sequences irrespective of whether they were slow-cooled (SC) or fast-cooled (FC) (Fig. 7E).

**Figure 7.**
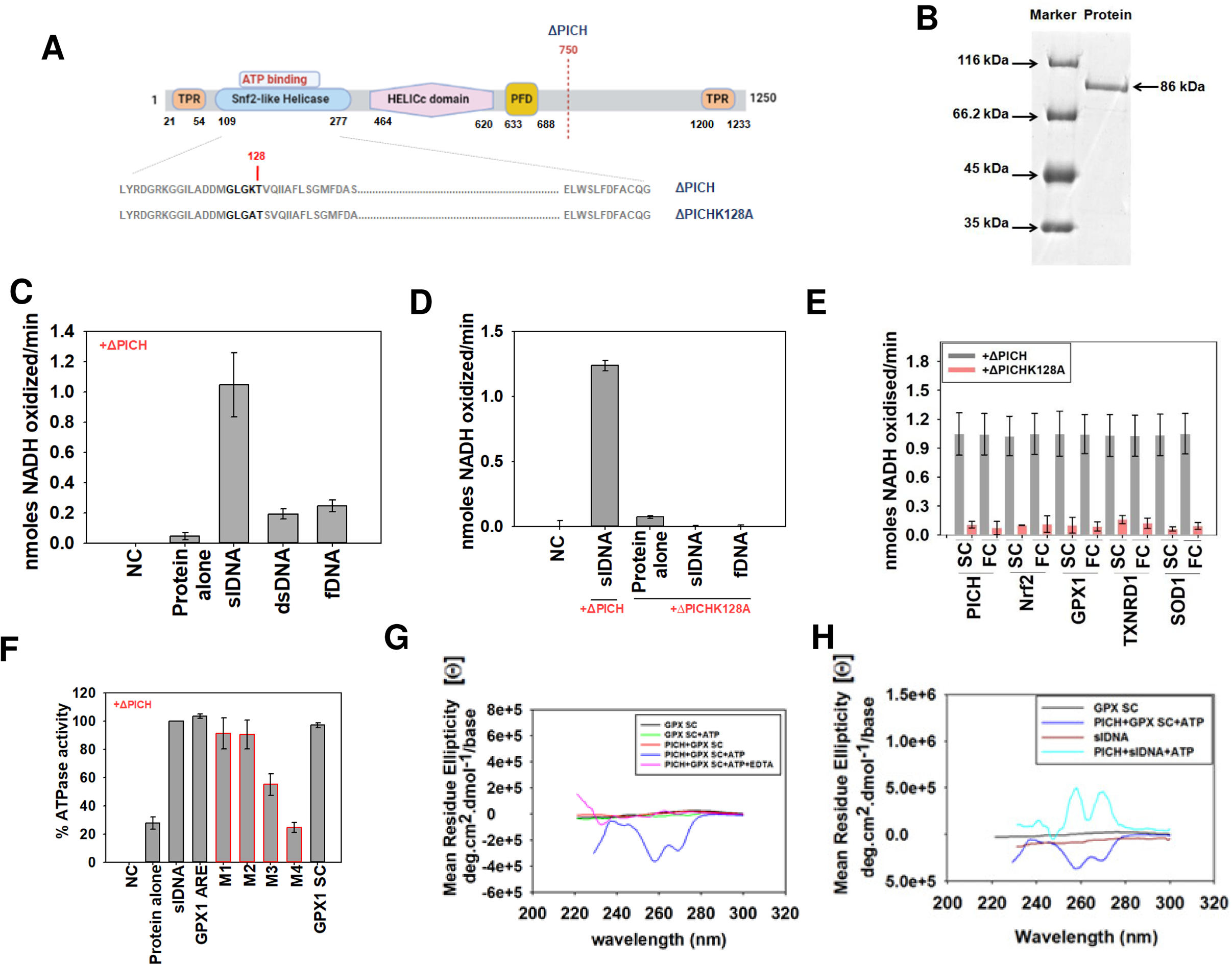
PICH alters the conformation of DNA in an ATP-dependent manner. (A). Representation of PICH domains. The cartoon shows the residue K128 which was mutated to alanine. (B). Coomassie Blue stained gel showing purified recombinant ΔPICH protein (86 kDa). (C). ATPase activity of ΔPICH protein was measured using different DNA effectors. (D). ATPase activity of the ΔPICHK128A and ΔPICH was measured in the presence of different DNA effectors. (E). ATPase activity ΔPICH was measured in the presence of DNA sequences where PICH binding was observed by ChIP. (F). ATPase assay of ΔPICH protein was measured in the presence of GPX1ARE wild type and mutated sequences. (G). CD spectra of slow-cooled GPX1SC DNA in the absence and presence of ΔPICH and ATP. (H). Comparison of the CD spectra of slDNA and GPX1SC in the absence and presence of ΔPICH and ATP. In the ATPase assays, a negative control (NC) that contained neither protein nor DNA was included.

To understand whether the sequence or the structure was the critical parameter in protein-DNA recognition, an oligonucleotide that represented only the ARE sequence of GPX1 promoter (GPX1 ARE) was designed (Supplementary Table 3). Mutations (M1 to M4) were introduced at selected positions (Supplementary Table 3). None of these oligonucleotides formed secondary structures (Supplementary Table 3). ATPase assays showed that the amount of ATP hydrolyzed was the same for both GPX1 ARE sequence and GPX1 SC (Fig. 7F). Next, we analyzed which of the residues are important for the interaction by mutation analysis. We found that when T at the 11^th^ position was mutated to A as in the case of M4 oligonucleotide, the ATPase activity was abolished suggesting that the protein bind to ARE in a sequence-specific manner (Fig. 7F).

To understand whether PICH can alter the conformation of DNA sequences, Circular dichroism (CD) spectroscopy was employed as studies have shown that different conformers of DNA can be identified by this methodology [35]. CD spectra of the GPX1SC DNA showed a large negative peak at 255 nm and a small negative peak around 270 nm in the presence of ATP and ΔPICH (Fig. 7G). The formation of these peaks was abrogated by the addition of EDTA indicating that the conformational change induced in the DNA by PICH was dependent on ATP hydrolysis (Fig. 7G). To understand whether PICH induced the same conformational change in all DNA molecules, the CD spectra of slDNA in the absence and presence of ATP and ΔPICH were also analyzed. In contrast to GPX1SC DNA, the slDNA formed two positive peaks at 252 nm and 265 nm, which was again dependent on the ATPase activity of the protein (Fig. 7H and Fig. S5B).

Thus, it was concluded that PICH possibly regulates the transcription of its effector genes by inducing conformational changes in the promoter regions.

## DISCUSSION

Oxidative stress reflects the imbalance between the expression of reactive oxygen species and the ability of the cell to detoxify or repair the damage through reactive intermediates. The reactive oxygen species cause DNA damage, lipid peroxidation, as well as oxidation of proteins. The cell combats the oxidative stress by producing antioxidants like superoxide dismutase, catalase, and glutathione peroxidase that can directly scavenge the free radicals. The expression of these antioxidant enzymes under transcriptional control wherein Nrf2, a transcription factor, has been shown to play a pivotal role. In addition, studies have shown that the expression of antioxidant genes is also co-regulated by chromatin remodeling mechanisms. KMT2D, a histone methyltransferase that catalyzes monomethylation of H3K4, has been shown to regulate the expression of antioxidant genes in prostate cancer [36]. Studies have also shown that transcriptional activation of *HO-1*, a gene encoding for homo oxygenase −1, by Nrf2 requires BRG1, an ATP-dependent chromatin remodeling protein [3]. However, there are no reports regarding the role of PICH in mediating transcriptional co-regulation of genes encoding for antioxidants during oxidative stress.

PICH was identified as a Plk1 kinase interacting protein and extensive studies have delineated its role in mitosis [5,37,38]. Further, studies have shown that PICH is an ATP-dependent translocase that can bind and translocate on double-strand DNA but cannot remodel nucleosomes [8]. Thus, the role of this protein in transcriptional co-regulation has not yet been investigated.

### PICH, a transcriptional regulator, manages the response to oxidative stress in HeLa cells

In this paper, the role of PICH as a transcriptional co-regulator when oxidative stress is generated in HeLa cells has been delineated. PICH regulates the expression of *Nrf2* as well as genes encoding for antioxidants in HeLa cells. Downregulation of PICH reduces the expression of Nrf2 as well as genes encoding for antioxidants leading to increased oxidative stress as well as DNA damage on treatment with H_2_O_2_. Overexpression of wild-type *PICH* in *PICH* depleted cells restored the expression of *Nrf2* and antioxidants in the presence of oxidative stress. The transcriptional regulation was dependent on the ATPase activity of PICH as overexpression of ATPase-dead mutant of *PICH* in *PICH* depleted cells failed to restore the expression of the effector genes. Finally, ChIP analysis confirmed that PICH can bind to the promoter regions of Nrf2 and antioxidant genes, thus directly regulating their expression.

The expression of *PICH* is regulated by Nrf2 in the presence of oxidative stress. *In silico* analysis identified ARE sequence in the promoter regions of *PICH* and ChIP experiments confirmed that Nrf2 occupancy increases on the *PICH* promoter only in the presence of oxidative stress. In contrast, PICH appears to regulate the expression of *Nrf2* both in the absence and presence of oxidative stress. Thus, PICH provides cellular defense by functioning as an oxidative stress induced response factor that regulates transcription of Nrf2 and antioxidant genes.

Analysis of the available ChIP-seq datasets showed that activating histone marks-H3K9ac, H3K4me2, H3K27ac, and H3K4me3- are enriched on *PICH*, *Nrf2*, *GPX1*, *TXNRD1*, and *SOD1* promoters. ChIP experiments confirmed that H3K27ac and H3K4me3 are enriched on the *PICH*, *GPX1*, *TXNRD1*, and *SOD1* promoter on oxidative stress while only H3K27ac is enriched on the *Nrf2* promoter. Thus, on oxidative stress, gene expression is regulated by Nrf2, a transcription factor, in collaboration with epigenetic regulators-PICH and activating histone modifications-leading to an open chromatin architecture and thus, increased transcription.

Depletion of *PICH* led to reduced enrichment of H3K27ac, RNAPII, and Nrf2 on the promoters of the responsive genes, suggesting that PICH is needed for the recruitment of the transcription machinery on oxidative stress. However, the order of recruitment cannot be delineated from these experiments as the expression of Nrf2 is affected in PICH depleted cells. There are multiple possibilities. One, PICH binds to ARE and recruits H3K27ac modifying enzyme along with RNAPII and Nrf2. The second possibility is that Nrf2 recruits PICH, histone modifying enzymes, and RNAPII. Finally, Nrf2 and PICH are both recruited by H3K27ac modification. The studies presented here do not allow us to differentiate between these different mechanisms because PICH depletion leads to downregulation of Nrf2. Additional experiments need to be performed to understand how these factors are recruited to the target genes on oxidative stress.

### PICH-Nrf2-H3K27ac complex possibly directs transcription output during oxidative stress

ChIP-reChIP experiments showed that PICH and Nrf2 are present simultaneously on the effector promoters. Co-immunoprecipitation showed that PICH and Nrf2 possibly form a complex both in the absence and presence of oxidative stress. Further, in the presence of oxidative stress, they appear to interact with the histone modification H3K27ac. Thus, the interaction between an ATP-dependent chromatin remodeler, a transcription factor, and histone modification might be driving the transcriptional output from effector genes on oxidative stress in HeLa cells.

### Mechanism of transcriptional regulation by PICH

As stated earlier, PICH does not appear to possess ATP-dependent nucleosome remodeling activity [8]. However, it can resolve triplex DNA [8]. BRG1, an ATP-dependent chromatin remodeling protein, is known to induce Z-DNA conformation on the *HO-1* promoter and this conformational change is associated with transcriptional activation of this gene on oxidative stress [3, 39]. SMARCAL1 too has been shown to alter the conformation of DNA present on the promoter regions in an ATP-dependent manner [20, 32]. Therefore, it was hypothesized that PICH might also alter the conformation of DNA in an ATP-dependent manner. CD spectroscopy showed that PICH can indeed alter the conformation of the DNA. The conformational change induced in the DNA appears to be dependent on the DNA sequence as slDNA and GPX1SCform different structures. The conformational change induced in a given DNA sequence is dependent on the ATPase activity of the protein.

Different DNA conformers have a varying degree of entanglement on a promoter region to make the transcription process feasible. Z-DNA, for example, is known to be stabilized transiently due to the generation of negative supercoiling by the movement of RNAPII, and thus, coupled to transcription regulation [40]. The CD spectra of the GPX1SC sequence show a large negative peak at 255 nm and another negative peak at 270 nm. These negative peaks are associated with a form of DNA known as X-DNA [35]. Movement of RNAPII across the DNA template generates positive supercoils this transcription-generated positive supercoiling is used to disrupt and/or eliminate road-block proteins, thus, destabilizing nucleosome structure making the underlying DNA more accessible to RNAPII [41, 42]. The X-DNA has been proposed to represent an overwound DNA that can potentially function as a sink for the positive supercoiling [43]. Thus, we can hypothesize that PICH possibly facilitates transcription by inducing an X-DNA structure that can act as an absorbent for positive supercoiling. However, further studies are needed to understand how the conformational change in the ARE sequences induced by PICH in the presence of ATP impacts transcription.

In conclusion, PICH appears to exert its effect through a combination of i) direct regulation of *Nrf2* expression; ii) ARE recognition and antioxidant response transcriptional regulation with Nrf2; iii) by interacting with histone mark H3K27ac as a part of active transcriptional machinery; and finally, iv) bringing chromatin conformational changes facilitating transcription, although the exact order of such events cannot be inferred from the study presented.

Oxidative stress is lethal to the cell and hence, the cell has devised mechanisms to alleviate it. PICH appears to be central to this process by regulating the expression of both antioxidant genes and Nrf2, the transcription factor involved in regulating the expression of antioxidant genes, thus expanding the role of this protein beyond mitosis into the realm of transcription regulation.

## MATERIALS AND METHODS

### Chemicals

DMEM, fetal bovine serum, antibiotic antimycotic solution, and trypsin-EDTA solution were purchased from Himedia (USA). Hoechst 33342 and Trizol were purchased from Sigma-Aldrich (USA). Hydrogen peroxide was purchased from Merck (Germany). RevertAid First Strand cDNA Synthesis Kit and TurboFect transfection Reagents were purchased from Thermo Fisher Scientific (USA). Restriction endonucleases were purchased from New England Biolabs (USA). SYBR Green PCR Master Mix was purchased from Kapa Biosystems (USA). Micro-amp Fast 96-well reaction plates, 0.1ml was purchased from Applied Biosystems (USA). QIAquick gel extraction kit was purchased from Qiagen (USA). Ni^+2^-NTA Sepharose resin was purchased from Novagen (Germany). Protein-G fast flow bead resin and Immobilon-P PVDF membrane were purchased from Merck-Millipore (USA). X-ray films, developer, and fixer were from Kodak (USA).

### Primers

Primers for ChIP, qPCR as well as oligonucleotides used for biophysical studies were synthesized either by Sigma-Aldrich (USA). or by GCC Biotech (India). The sequence of the primers and oligonucleotides used in this study is provided in Supplementary Tables 1-3.

### Antibodies

BRG1 (catalog#Ab70558), Histone H3 (catalog# Ab1791), RNAPII (Rpb1 CTD catalog# 2629), Nrf2 (catalog#Ab62352), PICH (catalog# Ab88560), H3K27ac (catalog#4729) antibodies were purchased from Abcam (USA). For western blot, Nrf2 (catalog# 16396-1-AP) was purchased from Proteintech Group (USA). H3K4me3 (catalog# C42D8) and IgG (catalog# 2729) were purchased from CST (USA). For immunocytochemistry, γH2AX (catalog# SAB5600038) was purchased from Sigma-Aldrich (USA). The secondary antibody (TRITC conjugated IgG and FITC conjugated IgG) was purchased from Genei, Bangalore, India.

### Transient downregulation of *PICH* and *Nrf2*

*PICH* and *Nrf2* were downregulated in HeLa cells using shRNA constructs against the 3′ UTR. A short region of 21 bases from the 3′ UTR was selected as the sense fragment for *PICH* shRNA (5′–CCGGTATTCTGAGCACTAGCTTAATCTCGAGATTAAGCTAGTGCTCAGAATATTT TTG-3′). Similarly, a short region from the 3′ UTR was selected as the sense fragment for *Nrf2* (5′-CCGGGCTCCTACTGTGATGTGAAATCTCGAGATTTCACATCACAGTAGGAGCTTTTTG-3′). These were ligated to the antisense fragment interrupted by a non-related spacer sequence. The double-stranded oligonucleotide encoding the shRNA was cloned into the pLK0.1 vector between AgeI and EcoRI restriction sites.

### Overexpression of *PICH* and *Nrf*2

The enhanced GFP tagged construct for *PICH*, pEGFP PICH (Nigg CB62) was a gift from Erich Nigg (Addgene plasmid # 41163; http://n2t.net/addgene:41163; RRID: Addgene_41163) [5].

pcDNA3-EGFP-C4-Nrf2 was a gift from Yue Xiong (Addgene plasmid # 21549; http://n2t.net/addgene:21549; RRID: Addgene_21549) [44]

### Cell culture and transfection

HeLa cells obtained from NCCS, Pune, India were cultured in DMEM containing 10% fetal bovine serum and 1X antibiotic antimycotic solution at 37°C and 5% CO_2_. For inducing oxidative stress, the cells were treated with 100 μM hydrogen peroxide for a specified time.

### RNA isolation and qPCR

Total RNA was extracted using the Trizol reagent. 90% confluent cells in a 35 mm plate were lysed with 1 ml of the Trizol reagent to give a homogenized lysate. The lysate was transferred to an Eppendorf tube. 200 μl of chloroform was added to each tube per ml of Trizol reagent, shaken vigorously, and allowed to stand for 10–15 min at room temperature. The samples were centrifuged at 11,000 rpm for 15 min at 4°C. The top aqueous layer was transferred to a fresh Eppendorf tube and 0.5 ml of isopropanol was added per ml of Trizol reagent, mixed, and allowed to stand at room temperature for 10–15 min. The samples were then centrifuged at 11,000 rpm for 10 min at 4°C. The RNA pellet obtained was washed with 70% ethanol and resuspended in DEPC-treated water. The concentration of the purified RNA was determined using NanoDrop 2000 (Thermo Fisher Scientific, USA) and an equal amount of RNA from various samples was used for preparing cDNA using random hexamer primers according to the manufacturer’s protocol. The prepared cDNA was checked for quality by performing a PCR using suitable primers.

Quantitative real-time RT-PCR (qPCR) was performed using the 7500 Fast Real-Time PCR system (ABI Biosystems, USA). Gene-specific primers designed for exon-exon junctions were used for qPCR. For each reaction,10 μl of samples were prepared in triplicates and the data obtained was analyzed using Fast7500 software provided by the manufacturer.

### Chromatin Immunoprecipitation (ChIP)

The cells were cross-linked for 10 min by adding formaldehyde (final concentration 1%) and later quenched by adding glycine (final concentration 125 mM) to the media. The cells were then washed thoroughly using ice-cold PBS and scraped into 1 ml buffer containing 150 mM NaCl, 0.02 M EDTA, 50 mM Tris-Cl (pH 7.5), 0.5% (v/v) NP-40 1% (v/v) Triton-X100 and 20 mM NaF. The cells were collected in Eppendorf tubes and pelleted at 12000g at 4°C for 2 min twice. The pelleted cells were treated with freshly prepared 150 µl of sonication buffer1 containing 0.01 M EDTA pH 8.0, 50 mM Tris-Cl (pH 8.0), and 1% SDS for 15 min at 4°C on a rocker. This was followed by addition of 50 µl of freshly prepared sonication buffer 2 containing 30 M EDTA 40 mM Tris-Cl (pH 8.0) 2% (v/v) NP-40, 0.04% (w/v) NaF. Sonication was done using a water bath sonicator (40 cycles of 30 sec pulse/ 20 sec rest). The sonicated samples were centrifuged, and the supernatant was used for further analysis. 50 μl of the sonicated sample was purified and DNA concentration was determined. An equal amount of chromatin (25 µg) was taken for performing IP using various antibodies. One sample was kept as Beads alone for negative control. Pre-adsorbed protein A bead resin (pre-adsorbed with 75 ng/μl sonicated salmon sperm DNA and 0.1 μg/μl of BSA) and 1.5 to 2 μg of the desired antibody was added to each sample and incubated overnight at 4°C on a rotator. This was followed by washing the pelleted bead resin two times in IP buffer containing 0.15 M NaCl, 0.02 M EDTA (pH 8.0), 50 mM Tris-Cl pH8.0,1% (v/v) NP-40,0.02% NaF,0.50% sodium deoxycholate and 0.1% SDS. This was followed by washing three times in wash buffer (0.5 M LiCl, 0.02 M EDTA (pH 8.0), 0.1 mM Tris-Cl (pH 8.0),1% (v/v) NP-40, 0.02% NaF and 1% sodium deoxycholate). The immune complexes were again washed twice in IP buffer and then a final wash was given in TE buffer (0.01 M Tris-Cl (pH 8.0), 0.001 M EDTA). The bound DNA was eluted using 100 µl of 10% (w/v) Chelex® 100 (Bio-Rad, USA) slurry prepared as per the manufacturer’s instructions. The eluted DNA was used for qPCR using standardized primers.

### Western blotting

Adherent cells in 100 mm dishes were washed with 1X PBS 3–4 times, scraped with a cell scraper, and collected in 1ml of 1X PBS. The cells were pelleted at 2500 rpm at 4°C. The cell lysate was prepared on ice using 300 μl of modified RIPA lysis buffer (50 mM Tris-Cl pH7.5, 150 mM NaCl, 2 mM EDTA, 0.5% sodium deoxycholate, 1% (v/v) NP-40, 0.1% sodium dodecyl sulphate, 1% TritonX-100 and 3 mM PMSF). The samples were sonicated for 3 cycles (20 sec on,40 sec off) using a sonicator. The protein concentration was determined using the Bradford assay. 100 µg protein was used to run the SDS-PAGE. Western blotting was done as described in (https://www.abcam.com/protocols/general-western-blot-protocol#loading-and-running-the-gel). The western blots were quantitated using Image J software.

### Oxidative stress assay

DCFDA (Thermo Fisher Scientific, USA) was prepared as a 10 mM stock in DMSO. Cells were seeded in a 35 mm dish to reach 40% to 50% confluency on the day of the experiment. The serum containing DMEM media was replaced with serum-free DMEM (SFM) media for 6 h before the start of the experiment to reduce the level of serum available in the medium.

At the time of the experiment, 10 µM DCFDA was added to the cells in dark and incubated at 37°C for 10 min. Subsequently, SFM containing 100 µM H_2_O_2_ was added, and the cells were further incubated for 20 min. Fluorescence was recorded for at least 100 cells using a fluorescent microscope (Eclipse Ti-E, Nikon) at the FITC range (495nm).

### Co-immunoprecipitation

HeLa cells were grown to 80% confluency. The cells were collected by trypsinization, and the cell pellet was resuspended in Buffer A (10 mM HEPES, (pH 7.4),10 mM KCl, 0.1 mM EDTA, 0.1 mM EGTA,1 mM DTT, and 0.5 mM PMSF) and incubated on ice for 15 min. Next, 6 µl NP-40 was added, vortexed for 15 sec, and the supernatant was collected by centrifuging the samples at 8000 rpm for 1 min. The supernatant was collected and kept on ice. The remaining pellet was re-suspended in 50 µl Buffer C (20 mM HEPES, (pH 7.4), 0.4 M NaCl, 0.1 mM EDTA,0.1 mM EGTA, 1mM DTT, and 0.5 mM PMSF). The samples were vortexed for 60 sec with 5 min intervals for the next 30 min. The supernatant was collected by centrifuging at 14000 rpm for 20 min and mixed with the previously collected supernatant. For immunoprecipitation, 600 μg of protein and 5-6 µg of specific antibody were mixed and incubated for 16 h at 4°C. The next day, 50 µl pre-equilibrated protein A bead resin was added and incubated on a rotating wheel for 3 h. Subsequently, the beads were collected by centrifugation at 2500 rpm for 5 min and washed twice with Buffer A. The samples were boiled with protein loading dye at 95°C for 10 min and centrifuged at 12000 rpm for 5 min at 4°C. The supernatant was collected in a fresh tube and loaded onto an 8 % SDS-PAGE. Western blotting was done as described in (https://www.abcam.com/protocols/general-western-blot-protocol#loading-and-running-the-gel). The western blots were quantitated using Image J software.

### Chip-reChip

ChIP-reChIP was performed as described in [45].

### Immunocytochemistry

Cells were grown on a coverslip in 35 mm dishes till 50% confluency. The cells were transfected with either ScrRNA or with the Sh*PICH* construct. After 36 h, the cells were either left untreated or treated with 100 µM H_2_O_2_ for 20 min at 37°C. The cells were then washed with PBS and fixed with 1:1 methanol: acetone for 20 min at 4°C. Subsequently, the cells were washed with 1X PBS and permeabilized with 0.5% of Triton X-100 in PBS for 20 min at 4°C. After washing with PBS, blocking was done with 5% BSA in PBS for at least 6 h at 37°C. Then the cells were incubated with primary antibody at 4°C overnight. The cells were then washed 5 times with 1X PBS and incubated with TRITC- and FITC-conjugated secondary antibody along with Hoechst stain for nuclear staining for 45 min at 37°C. After washing 6 times of 5 minutes each with ice-cold 1X PBS, the cells were fixed in 70% glycerol on a glass slide, sealed, and visualized under Nikon Eclipse TiE microscope. The images for at least 100 cells were analyzed using NiS v 4.0 software. In all the immunocytochemistry experiments 1:100 dilution of PICH antibody, 1:1000 dilution of γH2AX antibody, and 1:1000 dilution of Hoechst stain were used.

### Site-directed mutagenesis

All the mutants described in the present study were generated by PCR amplification using specific primers (Supplementary Table 4) and the mutations were confirmed by sequencing.

### Protein Purification

*E. coli* BL21 (DE3) was transformed with the plasmids expressing either the wild type or the K128A mutant. Protein expression was induced by adding 0.5 mM IPTG at 16°C for 16 h. The cells were harvested by centrifugation at 5000 rpm at 4°C and resuspended in lysis buffer containing 50 mM Tris-Cl (pH 8.0), 150 mM NaCl, 150 mM MgCl_2_, 0.1% (v/v) Triton X-114, 0.2 mg/ml lysozyme, 10 mM β-mercaptoethanol, and 0.5 mM PMSF. The cells were incubated at 4°C for 1 h and lysed by sonication (15 sec ON and 45 sec OFF; five cycles). After sonication, the cell debris was removed by centrifugation at 10000 rpm for 45 min at 4°C. The supernatant, thus, obtained was loaded onto a Ni^2+^-NTA column. The column was washed with wash buffer (50 mM Tris-Cl (pH 8.0), 150 mM NaCl, 150 mM MgCl_2_, 10 mM β-mercaptoethanol, 30 mM imidazole, and 0.5 mM PMSF) and the bound protein was eluted with buffer containing 100 mM imidazole, 50 mM Tris-Cl (pH 8.0), 5 mM MgCl_2_, 100 mM NaCl, and 5 mM β-mercaptoethanol. The purified fractions thus obtained were analyzed by 10% SDS-PAGE. The concentration of the purified protein was determined using the Bradford reagent.

### ATPase assay

NADH coupled oxidation assay was used to monitor the ATPase activity of purified PICH. Briefly, 0.2 μM of protein was incubated in 1X REG buffer (125 mM Tris-acetate (pH 8.0), 30 mM magnesium acetate, 300 mM potassium acetate, 25 mM β-mercaptoethanol, 6.8 mg/ml of phosphoenolpyruvate, 50 units of pyruvate kinase, and 50 units of lactate dehydrogenase) along with 20 nM DNA and 2 mM ATP, 0.1 mg/ml NADH containing in a 250 μl of reaction volume. The reaction was incubated for 15 min at 30°C and the absorbance was measured at 340 nm using a microplate spectrophotometer (BioTek®Synergy).

The sequences of DNA used in the ATPase assays are given in Supplementary Table 3.

### Catalase activity assay

The cell lysate was prepared on ice using 300 μl of modified RIPA lysis buffer (50 mM Tris-Cl (pH 7.5), 150 mM NaCl, 2 mM EDTA, 0.5% sodium deoxycholate, 1% (v/v) NP-40, 0.5% sodium dodecyl sulphate, 1% TritonX-100 and 3 mM PMSF). Subsequently, 0.95 ml of 0.1 M phosphate buffer (pH 7.4), 1.0 ml of freshly prepared 0.05 M hydrogen peroxide, and 100 μg supernatant were added in a 3 ml cuvette.

The optical density was read at 240 nm using Cary 60 UV-Vis spectrophotometer (Agilent Technologies, USA). The activity was calculated as follows:

Catalase activity (nmoles of H_2_O_2_ consumed/min/mg protein) = ΔOD/min x volume of assay x 10^9^ / MEC x volume of enzyme x mg protein x PL x VCF

Where:

Volume of assay = 3.0 ml MEC = 39.6 M^−1^ cm^−1^

Volume of enzyme = 0.05 ml

VCE = Volume conversion factor = 100

PL= path length = 1 cm

### Circular dichroism (CD)

To investigate the conformational changes in the DNA molecules, reactions were performed in 1 mM sodium phosphate buffer (pH 7.0). 500 nM DNA was incubated with 1 µM PICH, 10 mM Mg^+2^, and 2 mM ATP. The CD spectra (200 300 nm) were recorded using a Chirascan CD spectrophotometer (Applied Photophysics). To investigate the role of ATP hydrolysis in the conformational change 50 mM EDTA was used to inhibit the ATPase activity.

4 scans were measured for each reaction. The buffer and ligand contributions were subtracted. The spectra, thus, obtained in millidegrees was converted to mean residue ellipticity by using the formula: [θ] = (S x mRw)/(10 cl).

S is the CD signal in millidegrees, c is the protein concentration in mg/ml, mRw is the mean residue mass and l is the path length in centimeters.

### Statistical analysis

All qPCR and ChIP experiments are reported as average ± standard error of mean (SEM) of three independent experiments unless otherwise specified. The statistical significance (p-value) was calculated using paired t-test available in Sigma Plot. The differences were considered significant at p <0.05.

### Original blots

The original blots are provided in Supplementary Fig. S6.

## ACKNOWLEDGEMENTS

The authors would like to thank the Central Instrumentation Facility, School of Life Sciences for fluorescence microscope, and Advanced Instrumentation Research Facility, JNU, for the CD spectrophotometer.

